# ChondroMap: A Spatially Resolved Framework for Quantifying the Mechanical Exposure Between Cartilage and Bone Surfaces in Knee Osteoarthritis

**DOI:** 10.64898/2026.11.25.767888

**Authors:** Ziang Jiang, Tingyuan Wang, Tongxi Hu, Lemeng Dai, Stephan Reichenbach, Harald Bonel, Tristan Maerz, William R. Taylor, Qiang Zhang

## Abstract

**Objective:** This study presents “ChondroMap”, a spatially resolved framework for quantifying articulating cartilage- and bone-based contact interactions, demonstrated across various knee osteoarthritis (OA) severities.

**Design:** Through probing the interactions between segmented high-resolution, spatially resolved cartilage/bone MRI scans and articulating kinematics from biplanar video-radiography (BVR), the ChondroMap framework aggregates cartilage-on-cartilage (CoC), cartilage-on-bone (CoB), and bone-on-bone (BoB) interactions to form a cumulative heatmap of mechanical exposure. To illustrate the framework, three participants representing mild, moderate, and severe knee OA underwent non-weight-bearing MRI to segment femoral and tibial bone and cartilage surfaces, followed by tracking BVR (30Hz) during level walking. ChondroMaps of registered kinematic and morphological data aggregated the articulating mechanical interactions throughout the activity cycle.

**Results:** Application of the ChondroMap framework to the 3 participants revealed articular interaction patterns that varied markedly with severity. In the mild case, cumulative CoC strain was confined to a compact lateral region with minimal CoB and no BoB contact. The moderate case showed the greatest coexistence of all three interaction modes, including femoral- and tibial-sided CoB interactions and substantial medial BoB contact (up to 40%) adjacent to cartilage-related strain. The severe case showed consolidation into broad medial BoB regions (up to 65%), with high strains confined to the margins of denuded areas.

**Conclusion:** ChondroMap presents a framework to capture the magnitude, duration, and spatial distribution of cartilage- and bone-based tibiofemoral interactions across complete activity cycles, offering a promising basis for mechanistic characterization of the articular environment in knee OA, particularly in advanced or heterogeneous degeneration.

## Introduction

Knee osteoarthritis (OA) is a complex joint disease characterized by multiple pathological features, including spatially heterogeneous pathological remodelling of articular cartilage and subchondral bone (1, 2). Given the complex geometry of the articulating tissues, the local articular mechanical environment is governed not only by instantaneous loading magnitude, but also by the duration and spatial distribution of tibiofemoral surface interactions across activity cycles (3). Peak-load measures capture only a limited snapshot and cannot reflect the cumulative mechanical exposure experienced by the articulating surfaces, warranting temporally and spatially resolved characterization of articular interactions in knee OA throughout complete activity cycles. Such assessments are needed to identify patient-specific patterns of disease that can inform more personalized treatments and targeted interventions.

Biplanar video-radiography (BVR) with high-resolution anatomical models enables accurate reconstruction of *in-vivo* tibiofemoral kinematics and geometry-based estimation of articular contact (3). Contact is typically estimated either from overlap of opposing cartilage surfaces (3) or subchondral bone (4), and each provide only a partial description of the articulating mechanical environment. Importantly, most analyses of joint mechanics assume the presence of opposing cartilage surfaces, and may fail to detect cartilage-on-bone (CoB) or bone-on-bone (BoB) contact interactions at regions of severe cartilage thinning and/or osteochondral defects (5). Conversely, bone-based measures alone cannot distinguish compression of preserved cartilage from interactions between exposed subchondral bones. A unified framework tracking contact across cartilage and bone surfaces is therefore required to characterize the full spectrum of tibiofemoral interactions, particularly in joints with heterogeneous cartilage remodelling.

Loss of load-mediating cartilage unleashes a cascade of pathological processes that perpetuate OA progression, such as sclerotic remodelling of bone, phenotypic and structural alterations of the meniscus, and chronic synovial inflammation. Cartilage loss is partially driven by increased friction between the articulating (CoB or even BoB) surfaces (6), along with increased shear stresses and localized loading. As a result, understanding opposing-surface morphology, together with location and for how long cartilage and bone surfaces are mechanically exposed during functional activities is critical for establishing the relationships between subject-specific joint kinematics, kinetics, and structural damage in OA. Existing *in-vivo* contact analyses have focused on cartilage-on-cartilage (CoC) contact location, area, or deformation, mostly at selected or peak timepoints (7, 8). These approaches are limited to generic information when cartilage defects affect either or both opposing surfaces. Quantifying where opposing subchondral surfaces are exposed to CoB or BoB conditions, and for what proportion of the gait cycle such conditions exist, may provide an objective measure of local mechanical exposure across complete activity cycles. To this aim, we extend the concept of cumulative CoC strain (9) into a framework for bi-directional evaluation of cumulative CoB strain (CCoBS). Furthermore, spatially resolved BoB contact *frequency* indicates contact location and regularity. Together, these outcomes bridge the gap between whole-body and tissue-level mechanical conditions, and enable a more comprehensive characterization of the spatiotemporal articular environment in knee OA at various stages and in the context of focal cartilage lesions.

Accordingly, this study demonstrates “ChondroMap”, a subject-specific, spatially resolved framework characterizing tibiofemoral interactions involving both cartilage and subchondral bone. By incorporating MRI-derived models with tBVR-based kinematic analyses, we demonstrate the quantification of contact frequency and cumulative strain outcomes, enabling sensitive characterisation of the spatial distribution, magnitude, and temporal persistence of articular interactions across heterogeneous joint morphologies. As a proof of concept, we applied this innovative framework across OA severity grades to demonstrate its feasibility for tracking articular remodelling and its potential to generate patient-specific information that may aid in the improvement of diagnostic strategies and the evaluation of intervention or therapeutic efficacy.

## Methods

### Articular contact behaviour characterization

The ChondroMap framework initially requires access to a supine (unloaded) subject-specific bone-MRI, or a combined CT and MRI scan, spatially registered (8). Segmentation of the imaging datasets then provides bone morphology and cartilage surfaces as point clouds and polyhedron meshes (Fig. 1a). Baseline tibial 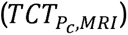 and femoral 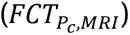 cartilage thicknesses were then determined at each cartilage cloud point, *P_c_*, as the shortest Euclidean distance between the MRI surface of the cartilage and the associated subchondral surface (6). By registering the skeletal geometry to dynamic tibiofemoral kinematics captured using BVR (Fig. 1b&c), ChondroMap then uses the point-cloud surfaces to distinguish different anatomically defined interaction modes: CoC, femoral or tibial CoB, and BoB.

**Figure 1.**
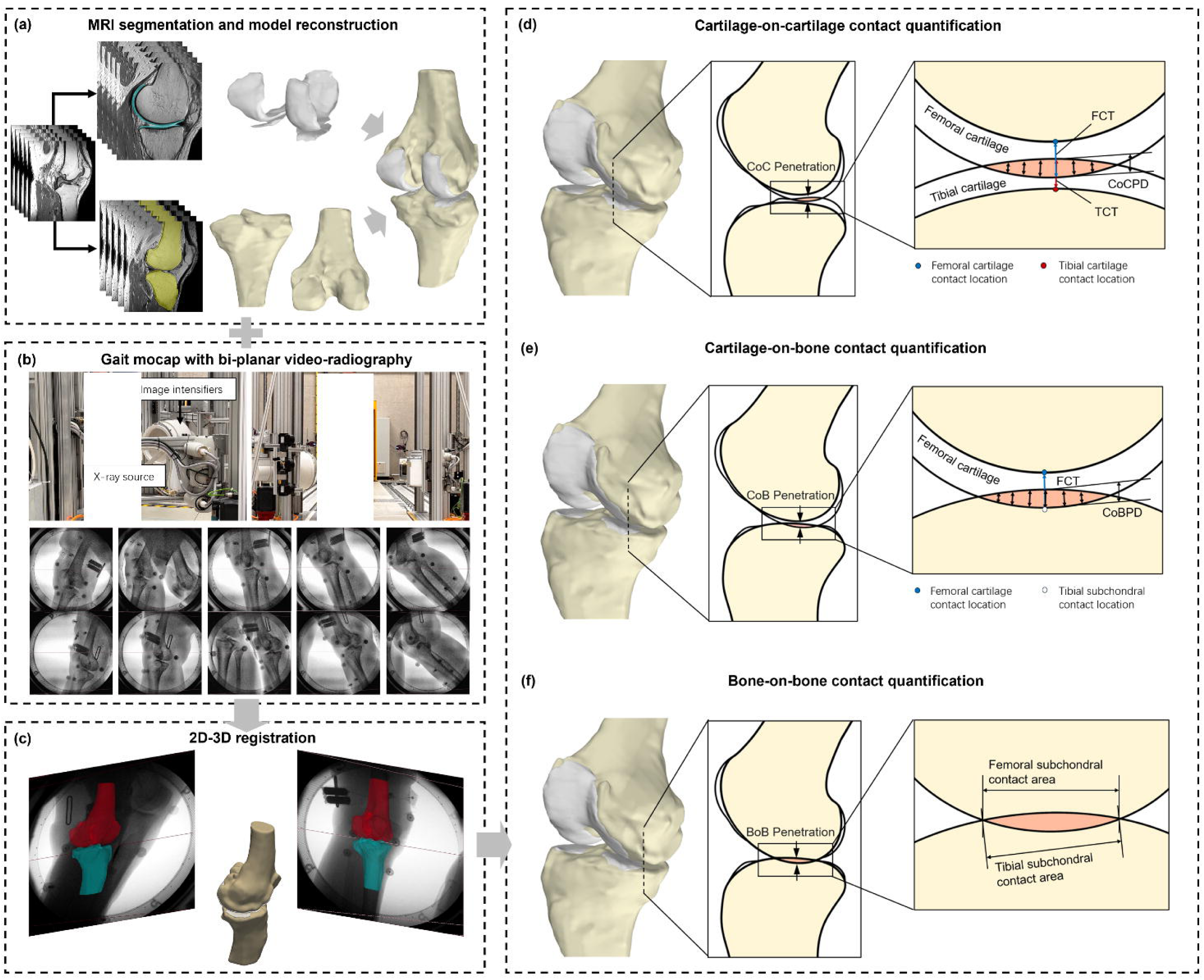
Overview of the ChondroMap workflow for imaging, kinematic reconstruction, and quantification of tibiofemoral cartilage and bone interactions. (a) Subject-specific femoral and tibial bone and cartilage surface models were reconstructed from MRI segmentations. (b) Tracking biplanar video-radiography (tBVR) captured knee motion during gait. (c) In-vivo tibiofemoral kinematics were reconstructed through 2D–3D registration of the bone models to the tBVR images. (d) For cartilage-on-cartilage interactions, local penetration depth (CoCPD) at each was normalized by the combined MRI-derived femoral and tibial cartilage thicknesses (FCT and TCT). The resulting strain was aggregated over the analysis period and normalized to its duration to derive cumulative cartilage strain. (e) For cartilage-on-bone interactions, local penetration depth (CoBPD) was normalized to the thickness of the existing cartilage and similarly aggregated and time-normalized to derive cumulative cartilage-on-bone strain. (f) Bone-on-bone contact was identified for occurrences of geometric inter-penetration between opposing subchondral bone surfaces at each frame, expressed as a frequency (% of the total frames analysed).

After skeletal registration to the kinematic (BVR) data, geometric penetration for local CoC interactions was quantified based on each *P_c_* (9). For every time-point (*t* ), a mesh-based point-in-polyhedron algorithm was used to determine whether each tibial *P_c_* penetrated the opposing femoral cartilage surface (Fig. 1d). In cases where penetration occurs, the CoC penetration depth 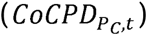 is calculated as the shortest overlapping Euclidean distance to the opposing cartilage surface. If no penetration was detected, a penetration depth of zero was assigned (Fig. 1d). To account for variations in the local cartilage thickness of each contact pair 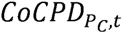 was normalized to the sum of 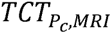 plus the associated instantaneous 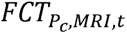, aggregated over the analysis period and normalized to cycle duration, *T* (from *t*1->*t*2), to derive the time-normalized cumulative CoC strain (*CCoCS_pc_*):

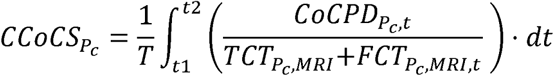

Similar pointwise algorithms were applied to calculate the CoB penetration depth (*CoBPD*) at each subchondral bone surface cloud point (*P_B_*), classified into two configurations according to whether the cartilage coverage was located on the femoral condyle 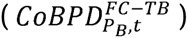 or the tibial plateau 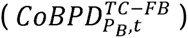 (Fig. 1e). Penetration depth was normalized by the corresponding MRI-derived thickness of the opposing cartilage only, and then aggregated over the analysis period and normalized to cycle duration, *T* , to derive the time-normalized cumulative cartilage strain:

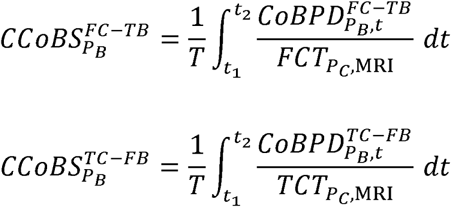

BoB contact frequency 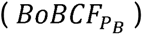 was determined based on geometrical penetration of the opposing bone surfaces as the aggregated sum of contact occurrences for each *P_B_* (Fig 1.f), expressed as a percentage of the total frames analysed:

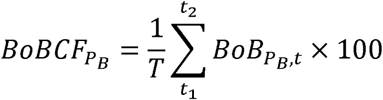

### Application of the ChondroMap Framework

This feasibility study was approved by the National Ethics Committee, and was conducted in accordance with the Declaration of Helsinki. Three participants (1 female and 2 males; mean age: 70.3yrs; mean BMI: 27.8 kg/m^2^), representing mild, moderate, and severe knee OA stages (Kellgren-Lawrence (K-L) grade II, III, and IV), were recruited. This study was conducted in accordance with the Declaration of Helsinki. Ethical approval was obtained from the Cantonal Ethics Committee of Zurich (BASEC 2025-01195), and all participants provided written informed consent prior to study participation.

### MRI and knee modelling

After 1h of non-weight-bearing, MRI scanning was performed (3-Tesla, Magnetom Vida, Siemens AG) with participants positioned supine and the knee fully extended. Proton density-weighted sequences were acquired in an oblique sagittal orientation with a 200×200mm field of view, 1mm slice thickness, and in-plane voxel size of 0.14×0.14mm. MRI images were segmented using automated thresholding and manual refinement (Mimics v28, Materialise AG), supervised by an experienced radiologist (H.B.). 3D surface models of the femur, tibia, and corresponding articular cartilage were reconstructed with an approximate mesh vertex spacing of 0.5mm.

### Video-radiography motion capture and data processing

Participants performed level walking tests at a self-selected speed, during which pulsed X-ray images (30Hz, resolution 1664 × 1600 pixels) of each participant’s index knee were acquired using tracking BVR. The calibrated X-ray images of each spatial pose, as well as segmented tibial and femoral bone models were imported into an in-house Python software for optimised 2D-3D registration, manually refined where necessary. The relatively 3D poses of the corresponding cartilage models were inherently preserved throughout the rigid-body transformations. To illustratively compare the mechanical exposure between mild, moderate, and severe cases of knee OA, the registered datasets of each subject were subsequently processed using the ChondroMap framework.

## Results

Segmentation of MRI data revealed varying cartilage thicknesses between the three participants, with distinct regions of denuded subchondral bone apparent, especially in the more severe cases (Fig. 2a). The ChondroMap framework revealed distinct spatial patterns of CoC, CoB, and BoB interactions throughout the measured gait activity (Fig. 2b). The contact interactions, as well as their distributions varied markedly across the illustrative mild, moderate, and severe cartilage loss cases. In the mild K-L II case, CoC interaction was concentrated within a compact region of the lateral femoral condyle and the corresponding lateral tibial plateau. CCoBS was restricted to a narrow region adjacent to the principal CCoCS distribution, while no BoB interaction was detected.

**Figure 2.**
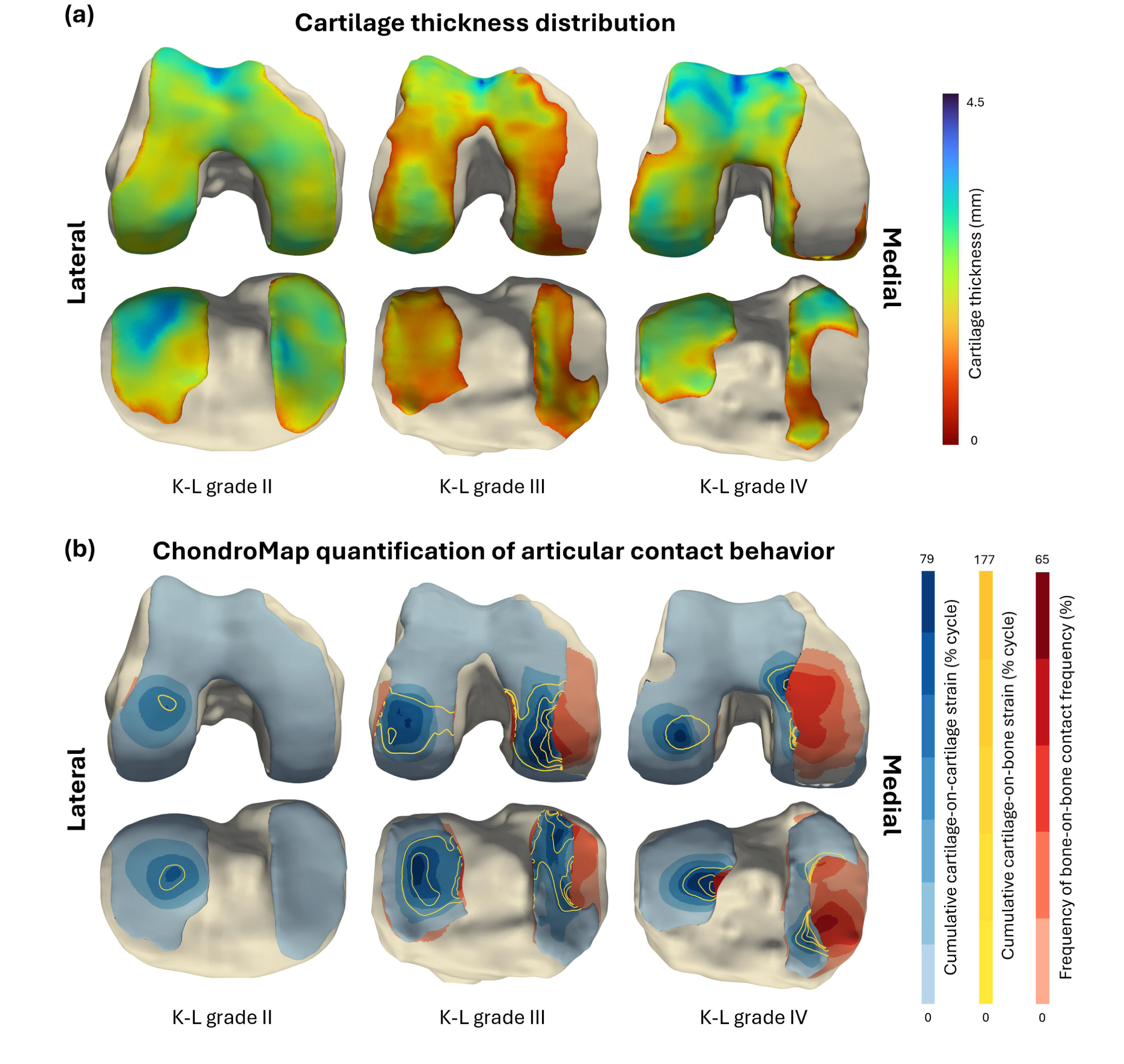
Participant-specific distributions of cartilage thickness and tibiofemoral ChondroMap outcomes during gait. Three illustrative cases representing Kellgren-Lawrence (K-L) grade II, III, and IV are shown from left to right. (a) MRI-derived cartilage thickness maps, together with uncoloured regions indicating the absence of cartilage and exposure of the underlying subchondral bone. (b) Time-normalized cumulative cartilage-on-cartilage strain (blue shading), cumulative cartilage-on-bone strain (CCoBS, yellow contours), and bone-on-bone contact frequency (BoBCF, red shading) are mapped onto the articular surfaces. CCoBS was quantified separately for both femoral-cartilage\tibial-bone interactions, and tibial-cartilage\femoral-bone interactions. BoBCF represents the percentage of analysed frames in which geometric bone-on-bone contact was detected.

The moderate K-L III case exhibited greatest spatial coexistence of the three interaction modes, with pronounced CCoCS distributions observed in both the medial and lateral compartments. Multiple CCoBS contours were identified on both the femoral and tibial surfaces, indicating the presence of TC–FB and FC–TB interactions. Within the medial compartment, cartilage-related cumulative strain was located adjacent to substantial and corresponding regions of BoB interaction on the femoral condyle and tibial plateau, where local contact frequency reached up to 40%. Smaller regions of BoB interaction were observed towards the peripheral lateral surfaces.

In the K-L IV case, focal CCoCS distributions remained evident in both the lateral and medial compartments, accompanied by localized CCoBS contours, which were concentrated along the margins of the denuded subchondral regions. In the medial compartment, broad BoB interaction regions occupied the opposing tibiofemoral subchondral surfaces exposed by extensive cartilage loss, and BoBCF reached up to 65%.

## Discussion

Understanding how tibiofemoral articular surfaces interact during functionally loaded activities is essential for quantifying patient-specific disease progression and treatment response (10). This is particularly relevant in advanced knee OA, where cartilage loss exposes the subchondral bone plate and severely compromises a principal load-attenuating interface (11). This study presented the ChondroMap framework for determining the extent to which such cartilage loss, coupled with pathological kinematics, leaves articular cartilage and subchondral surfaces mechanically endangered during gait. This approach lays the foundations for improved understanding of how mechanical exposure to the joint relates to disease severity and pathological tissue remodelling.

Comparison with the MRI-derived thickness maps showed close spatial correspondence between the ChondroMap outputs and cartilage morphology. In the mild case, coverage remained largely continuous, consistent with the absence of BoB interaction. Contours of CoB contact nonetheless appeared in this region, indicating apparent overlap between cartilage and the opposing bone surface for individual time frames. This overlap is most plausibly an artefact of the tBVR 2D 3D registration process, which introduces kinematic inaccuracy into the analysis. As registration accuracy for this challenging aspect of motion analysis improves, such artefacts should be reduced and eventually eliminated from ChondroMap analyses.

In the moderate case, pronounced CCoCS and CCoBS coincided with reduced or heterogeneous cartilage, most evidently near the medial defect margins. Here, all three modes co-existed, and cartilage strain lay directly adjacent to substantial BoB regions, which is interpretable as a transitional state in which residual marginal cartilage still mediates and redistributes load while the adjacent denuded bone already bears the joint contact forces directly. In the severe case, CCoBS was concentrated specifically along the denuded margins. This is consistent with reports that contact stresses and strains are elevated adjacent to defect rims, scaling with defect size (12). These ChondroMaps are therefore highly consistent with previous studies suggesting denuded bone reflects lost load carrying ability (2), and that bone marrow lesions colocalise almost exclusively with denuded cartilage (1).

In comparison with the mild OA case, it is observed that the severe case did not simply intensify a single loading mode but altered how load was discretised across interaction types. BoBCF rose from almost completely absent (mild) to 40% (moderate), and up to 65% (severe), but this rise came with a qualitative shift: the moderate case showed the greatest dispersion and coexistence of all three modes, while the severe case showed consolidation into broad, contiguous medial BoB regions with CCoCS/CCoBS confined to the margins. Future large-cohort studies using the ChondroMap approach are warranted to investigate the generalization of these specific patterns in knee OA at various K-L grades. Nevertheless, this shift toward larger, more conforming osseous contact is consistent with the recruitment of less functional cartilage regions and with known differences in tibiofemoral contact area between OA and healthy knees under load (13). Gait adaptations typical of advancing pathology (reduced knee loading demand, altered muscular contraction patterns, and slower gait speeds etc.) may further shape where and how these articular interactions occur (14, 15), even though potential kinematic modification patterns were not analysed in this study.

It should be emphasised that these candidate explanations and the underlying severity patterns were derived from three representative participants used to illustrate the ChondroMap framework itself, rather than from a cohort sufficiently powered to test severity effects. Larger, statistically powered studies across a broader severity spectrum are required to determine whether the patterns of mechanical exposure observed here generalise to larger populations and to clinical observations. Moreover, the ChondroMap framework does not yet incorporate cartilage’s viscoelastic or frictional properties since articular interactions were defined geometrically as mesh interpenetration rather than mechanical stress. An additional limitation of this study is that meniscus was excluded from the framework, as accurate segmentation of meniscus in OA MRI is technically difficult. Future work should incorporate meniscus into the ChondroMap and investigate its contact interactions with other tissues. Finally, the relationships between mechanical exposure assessed using ChondroMap and cartilage thickness have only been described qualitatively; future work should investigate any correlations using formal co-localization metrics.

In conclusion, this proof-of-concept study demonstrates ChondroMap, a unified, spatially resolved framework for characterizing dynamic tibiofemoral interactions in a subject-specific manner across articular cartilage and subchondral bone. By capturing the magnitude, duration, and location of cartilage- and bone-based interactions throughout complete activity cycles, ChondroMap thereby presents a viable framework for understanding the mechanical loading conditions to which articulating surfaces are exposed, particularly in joints with heterogeneous and advanced degeneration.

## Author Contributions

Conception and design: Z.J., W.R.T., and Q.Z.

Analysis and interpretation of the data: Z.J., T.W., T.H., and L.D.

Drafting of the article: Z.J. and Q.Z.

Critical revision of the article for important intellectual content: S.R., W.R.T., and Q.Z.

Final approval of the article: All authors.

Provision of study materials or patients: S.R. Statistical expertise: Q.Z.

Logistic support: S.R., H.B., and T.M.

## AI Use Disclosure

The authors declare that no generative AI tools were used in the preparation of this manuscript.

## Data Availability

The raw data generated and analysed in this study are not publicly available because they contain information that could compromise the privacy of the research participants, and public sharing is not permitted under the ethics approval (Cantonal Ethics Committee Zurich, BASEC 2025-01195) and the participants’ informed consent. De-identified data are available from the corresponding author upon reasonable request for non-commercial academic research. Requests will be answered within four weeks. Data access requires a signed data transfer agreement with ETH Zurich and, where required, approval from the responsible ethics committee. The data may be used only for the purpose specified in the request and may not be shared with third parties or used to re-identify participants.

## Reference

1. Bowes MA, McLure SW, Wolstenholme CB, Vincent GR, Williams S, Grainger A, et al. Osteoarthritic bone marrow lesions almost exclusively colocate with denuded cartilage: a 3D study using data from the Osteoarthritis Initiative. Ann Rheum Dis. 2016;75(10):1852–7.

2. Frobell RB, Wirth W, Nevitt M, Wyman BT, Benichou O, Dreher D, et al. Presence, location, type and size of denuded areas of subchondral bone in the knee as a function of radiographic stage of OA - data from the OA initiative. Osteoarthritis Cartilage. 2010;18(5):668–76.

3. Hosseini A, Van de Velde SK, Kozanek M, Gill TJ, Grodzinsky AJ, Rubash HE, et al. In-vivo time-dependent articular cartilage contact behavior of the tibiofemoral joint. Osteoarthritis Cartilage. 2010;18(7):909–16.

4. Dessinger GM, LaCour MT, Dennis DA, Kleeman-Forsthuber LT, Komistek RD. Can an OA Knee Brace Effectively Offload the Medial Condyle? An In Vivo Fluoroscopic Study. J Arthroplasty. 2021;36(4):1455–61.

5. Wirth W, Ladel C, Maschek S, Wisser A, Eckstein F, Roemer F. Quantitative measurement of cartilage morphology in osteoarthritis: current knowledge and future directions. Skeletal Radiol. 2023;52(11):2107–22.

6. Sutter EG, Widmyer MR, Utturkar GM, Spritzer CE, Garrett WE, Jr., DeFrate LE. In vivo measurement of localized tibiofemoral cartilage strains in response to dynamic activity. Am J Sports Med. 2015;43(2):370–6.

7. Mayr MF, Meine H, Lange T, Yilmaz T, Taghizadeh E, Schmal H, et al. In vivo kinematics of knee joint cartilage and meniscus contact areas under load application: a biomechanical MRI study. Arch Orthop Trauma Surg. 2026;146(1).

8. Xu C, Aloi N, Gale T, Nishida K, Fu F, Anderst W. Symmetry in knee arthrokinematics in healthy collegiate athletes during fast running and drop jump revealed through dynamic biplane radiography. Osteoarthritis Cartilage. 2023;31(11):1501–14.

9. Jiang Z. WT, Hu T., Dai L., Hosseini Nasab S.H., Reichenbach S., Bonel H., Maerz T., Zhang Q., Taylor W.R. A Novel Approach for Evaluating Cumulative Cartilage Compression and Strain. Osteoarthritis Cartilage. Under review.

10. Amiri P, Davis EM, Outerleys J, Miller RH, Brandon S, Astephen Wilson JL. High tibiofemoral contact and muscle forces during gait are associated with radiographic knee OA progression over 3 years. Knee. 2023;41:245–56.

11. Moisio K, Eckstein F, Chmiel JS, Guermazi A, Prasad P, Almagor O, et al. Denuded subchondral bone and knee pain in persons with knee osteoarthritis. Arthritis Rheum. 2009;60(12):3703–10.

12. Gratz KR, Wong BL, Bae WC, Sah RL. The effects of focal articular defects on cartilage contact mechanics. J Orthop Res. 2009;27(5):584–92.

13. Shin CS, Souza RB, Kumar D, Link TM, Wyman BT, Majumdar S. In vivo tibiofemoral cartilage-to-cartilage contact area of females with medial osteoarthritis under acute loading using MRI. J Magn Reson Imaging. 2011;34(6):1405–13.

14. Zeni JA, Higginson JS. Knee osteoarthritis affects the distribution of joint moments during gait. Knee. 2011;18(3):156–9.

15. Aydemir B, Huang CH, Foucher KC. Gait speed and kinesiophobia explain physical activity level in adults with osteoarthritis: A cross-sectional study. J Orthop Res. 2023;41(12):2629–37.

